# Phylogeny-aided detection of contamination in nearly 5 million SARS-CoV-2 genomes

**DOI:** 10.64898/2026.08.07.743473

**Authors:** Olivier Anoufa, Nhan Ly-Trong, Nick Goldman, Nicola De Maio

## Abstract

Contamination can occur during genome sequencing when a contaminant genome is accidentally mixed with the intended genome to be sequenced. Contamination can lead to incorrect consensus genome calling, disrupting analyses of pathogen evolution and transmission. To investigate the extent of this issue, we developed PhyCD, a phylogeny-aided computational approach to investigate contamination in SARS-CoV-2 genome sequencing data. PhyCD masks consensus genome positions associated with suspicious sequencing read coverage drops, then leverages pandemic-scale phylogenetic placement techniques to identify putative contamination events. Applying PhyCD to nearly 5 million SARS-CoV-2 genomes, we identified 10,942 putative contamination events under conservative parameters. Across the flagged genomes, PhyCD flagged — and so permits masking of — a total of 64,753 substitutions that could cause errors in downstream genome data analyses.

## 1 Introduction

In a global effort to respond to the COVID-19 pandemic, by early 2022 around 7 million SARS-CoV-2 genomes had already been sequenced and publicly shared. NCBI [25] and other public repositories now host over 9 million unrestricted open access SARS-CoV-2 genomes, while GISAID [26] hosts more than 17 million genomes under a controlled-access model. Remarkably, SARS-CoV-2 alone accounts for over 20% of all genomes ever sequenced, underscoring the immense scale of data generated during the pandemic [11]. This scale allows detailed reconstruction of pathogen evolution and spread [15, 17, 35], but it also introduces significant challenges. In particular, the need to process and analyse this vast amount of data in limited time increases the chance of phylogenetic tree reconstruction errors, sequencing artefacts, and other inconsistencies [4, 7, 11, 29]. Contamination is a major cause of sequence artefacts [18, 24, 32, 34]. Hence, methods tackling contamination issues could help prevent many issues in downstream analyses and therefore improve genomic surveillance accuracy and pandemic preparedness [10].

Contamination happens when multiple genomes are accidentally mixed together within one sequenced sample (Figure 1A). We focus here on contamination caused by multiple SARS-CoV-2 genomes from different hosts within a sample, which is harder to detect and account for than contamination from different species. Typically, one genome in a contaminated sample has substantially higher abundance than the other(s), and this is thought to be usually the genome of the case being originally sampled. This means that, in the absence of other issues, the consensus sequence of the sample would likely match this majority genome. Since only the consensus genome of a sample is usually used in downstream analyses, this type of contamination should not typically lead to biases and artefacts.

**Figure 1:**
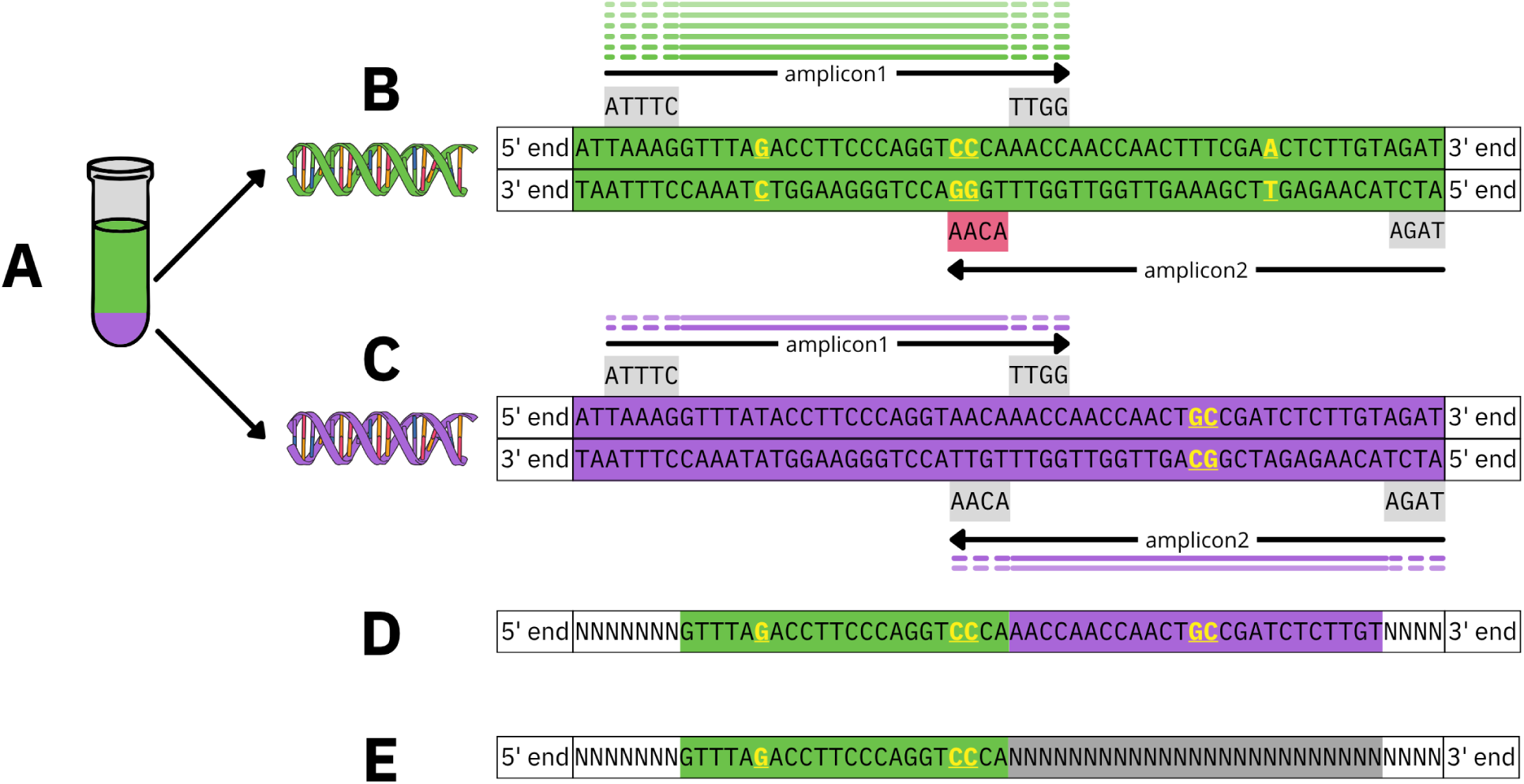
Sequencing of a contaminated sample. **A,** The contaminated test tube: in green is the de-sired majority genome, while in purple is the minority contaminant. **B,** Majority genome, containing substitutions T to G, AA to CC and T to A (coloured yellow). Two amplicons (black arrows) and their primers are highlighted. Amplicon1 is amplified normally, creating a large number of reads (greenlines) owing to the majority genome’s high abundance and correctly capturing the T to G and AA to CC substitutions. However, the AA to CC substitution is at the 3 primer binding site of amplicon2 (red box), leading to primer binding failure and therefore to amplicon2 dropout. **C,** Contaminant minority genome. This genome contains a TT to GC mutation within the amplicon2 region, but not affecting primer binding, and lacks the AA to CC primer-site mutation. Therefore both amplicons are amplified normally, although in low numbers of reads (purple lines) owing to the low abundance of contaminant in the sample. **D,** Consensus sequence obtained from this sample. On amplicon1, the reads of the majority genome are more abundant, so they inform the consensus sequence, which there-fore contains the T to G and AA to CC substitutions. On amplicon2, the reads from the contaminant genome are more abundant, so they inform the consensus sequence (which therefore contains the TT to GC substitution that is absent from the majority genome, and lacks the actual T to A substitution in the majority genome). **E,** Ideal outcome of our pipeline, where the genome regions informed by the contaminating genome are masked from the consensus sequence. This would capture the majority genome’s T to G and AA to CC mutations and omit the contaminant’s TT to GC mutation. We do not attempt to infer the majority genome’s unobserved T to A mutation. In **D** and **E,** note that primer sequences are usually trimmed from read ends; hence they do not inform the consensus sequence.

However, with amplicon sequencing (which has been adopted for sequencing the vast majority of SARS-CoV-2 genomes), mutations at genome positions of amplicon primers can cause these primers not to bind to the genome, and so the corresponding amplicon not to be sequenced (“amplicon dropout”: see Figure 1B and [2, 9, 22, 24]). Primers are short tailored sequences of DNA that bind to a specific location of the genome. During PCR amplification, they provide a starting and ending point to the polymerase. The amplified region between two primers is called an amplicon and sequences generated by replicating this region are called reads. Many of the primer binding issues can be addressed with regular updates to primer schemes, but substantial numbers of samples are still affected if sequenced before the relevant primer scheme update [3, 11, 12, 30, 31]. If amplicon dropout occurs in the majority genome but not a minority one within a contaminated sample (Figure 1B,C) then sequencing reads from the minority contaminating genome can be the only ones covering part of the genome. The consensus sequence for these positions called from the read data of this sample will then reflect the contaminating minority genome, rather than the majority one, resulting in a hybrid consensus of the two genomes resembling recombination [24] (Figure 1D). This can be problematic because the minority genome might contain at these positions mutations not present in the majority genome, resulting in artefactually inferred mutations, lineages, and recombination events in downstream analyses [2, 24].

Some authors have proposed *in vitro* solutions to address contamination in SARS-CoV-2 genomes, for instance spike-in barcoding [14]. Others have instead proposed *in silico* methods to address contamination or phenomena that, at least in the read data, cause similar patterns such as mixed infections or long-term within-host evolution. Eyre et al. [8] for example proposed a likelihood-based model of mixed bacterial infections applied to read data nucleotide counts. Sobkowiak et al. [27] proposed a Bayesian model-based clustering for the same task, applying a Gaussian mixture model to the allele frequencies of Single-Nucleotide Polymorphisms (SNPs) at heterozygous sites. Töpfer et al. [28] presented QuasiRecomb, a Hidden Markov Model (HMM) to infer different within-host viral haplotypes and their recombinants. Wagatsuma et al. [33] developed the tool vClean to tackle contamination in viral metagenomics; vClean uses machine learning models to estimate contamination probabilities. Krasilnikova et al. [13] proposed Polyphonia, a method aimed at detecting inter-sample contamination from sequencing data; their model considers a pair of samples at a time (a putative contaminated sample and a putative contaminant) and checks if the mutations of the contaminant are present at a consistent minor proportion in the potentially contaminated sample. Finally, Pipek et al. [19] investigated co-infection and intra-host recombination in more than 2 million samples; the method leverages lineage-defining mutations to detect samples that have SNPs corresponding to different strains. While all these approaches address (or might be used to address) contamination, none accounts for the effect of amplicon dropouts, which can dramatically alter the relative abundance of the two genomes in the sample (and therefore the consensus sequence) and which is our main focus.

We present PhyCD (Phylogeny-aided Contamination Detection), an approach to detect and address the impact of contamination (and related phenomena like mixed infections) in SARS-CoV-2 genome data. Our approach:

- scales efficiently to millions of SARS-CoV-2 sequencing datasets;
- integrates multiple sources of information including variation in coverage along the sequenced genome, within-sample heterozygosity, and a massive reference phylogeny;
- explicitly targets amplicon dropouts to address their detrimental effect on consensus genome calls, even at very low levels of contamination that might not cause detectable within-sample heterozygosity.

We apply PhyCD to nearly 5 million SARS-CoV-2 genomes and show that phylogenetic information can provide reliable signal for contamination detection at pandemic scale, revealing that around 0.1% of the investigated genomes show signs of contamination.

## 2 Methods

### 2.1 Data

We analyse a public dataset of 4,952,451 SARS-CoV-2 genomes assembled using Viridian [11], a tool that addresses several systematic issues in SARS-CoV-2 sequencing data processing, including some pervasive reference biases. Viridian achieves this by:

- masking consensus genome positions with low coverage (<20X) rather than using a reference genome to fill the gaps in the assembly;
- inferring the primer scheme being used in the considered sample and consequently trimming primer sequences within reads;
- masking consensus genome positions with substantial heterozygosity within the considered sample, which should prevent many potential consensus calling issues caused by contamination with a high-abundance contaminant.

We further mask 47 genome positions affected by recurrent consensus sequence errors [7] and the first 72 and last 134 genome positions that are often not reliable [4].

However, these steps do not address issues caused by a combination of low-concentration contamination and amplicon dropout (Figure 1), specifically in case the minority contaminant reads successfully amplify at the dropout site with read depth above 20X, in which case we expect Viridian (and similarly most assemblers) to call the contaminant alleles as the consensus at these positions. We address this in PhyCD by targeting both relative drops in read depth and absolute low depth. Our pipeline, described below, identifies and adjusts consensus sequences potentially affected by these issues.

### 2.2 Dropout masking

Our objective is to mask portions of a consensus sequence in case they might have been affected by contamination and amplicon dropout, where the coverage of the main genome in a hypothetical contamination might have dropped to almost 0 and reads of a potential contaminant might have become relatively more prevalent. We aim to achieve this using a consensus sequence masking approach that we call “dropout masking”, based on the level of read coverage in a sample compared to the median coverage in the same sample. The idea is that in the presence of an amplicon dropout and contamination we expect no reads from the main genome at the considered amplicon, and only reads from the (low-abundance) contaminant, therefore resulting in a considerable drop in coverage at the amplicon. Specifically, we use a threshold *ρ*, and in a sample *s* with median coverage *M_s_* we mask positions with coverage below *ρM_s_*. We also impose a minimum coverage threshold of 50X to prevent unreliable consensus calls from positions with low coverage, so overall we mask any position with coverage below max(50, *ρM_s_*).

We want to use a balanced value of *ρ*, since too stringent a threshold can cause excessive loss of consensus sequence resolution while an over-lenient one might leave potential consensus sequence errors unmasked. We tested four different *ρ* values: 5%, 10%, 15% and 20%, with results discussed in Section 3.1. Note that, because Viridian subsamples reads up to a maximum coverage of around 1,000–2,000 at each position, our approach cannot mask potential amplicon dropouts that would result in a final coverage above around 1000*ρ*–2000*ρ*.

### 2.3 Our pipeline

We consider the following evidence of contamination within the samples considered:

- multiple heterozygous genome sites at consistent minor allele frequency proportion, as also considered by other similar methods [8, 13]; on its own, however, this information might not be sufficient to detect contamination at very low proportion, which might still impact consensus calling due to amplicon dropout or high variation in allele proportions at heterozygous sites [1].
- substitutions at regions of high coverage drops compatible with a combination of contamination and amplicon dropout.

We primarily focus on the second form of evidence given that it is the one most likely to reveal consensus sequence errors. We consider heterozygous sites only in the late stages of the pipeline to corroborate putative contaminated samples. Our pipeline, “PhyCD”, is summarized in Figure 2 and each of its steps is described in detail below.

**Figure 2:**
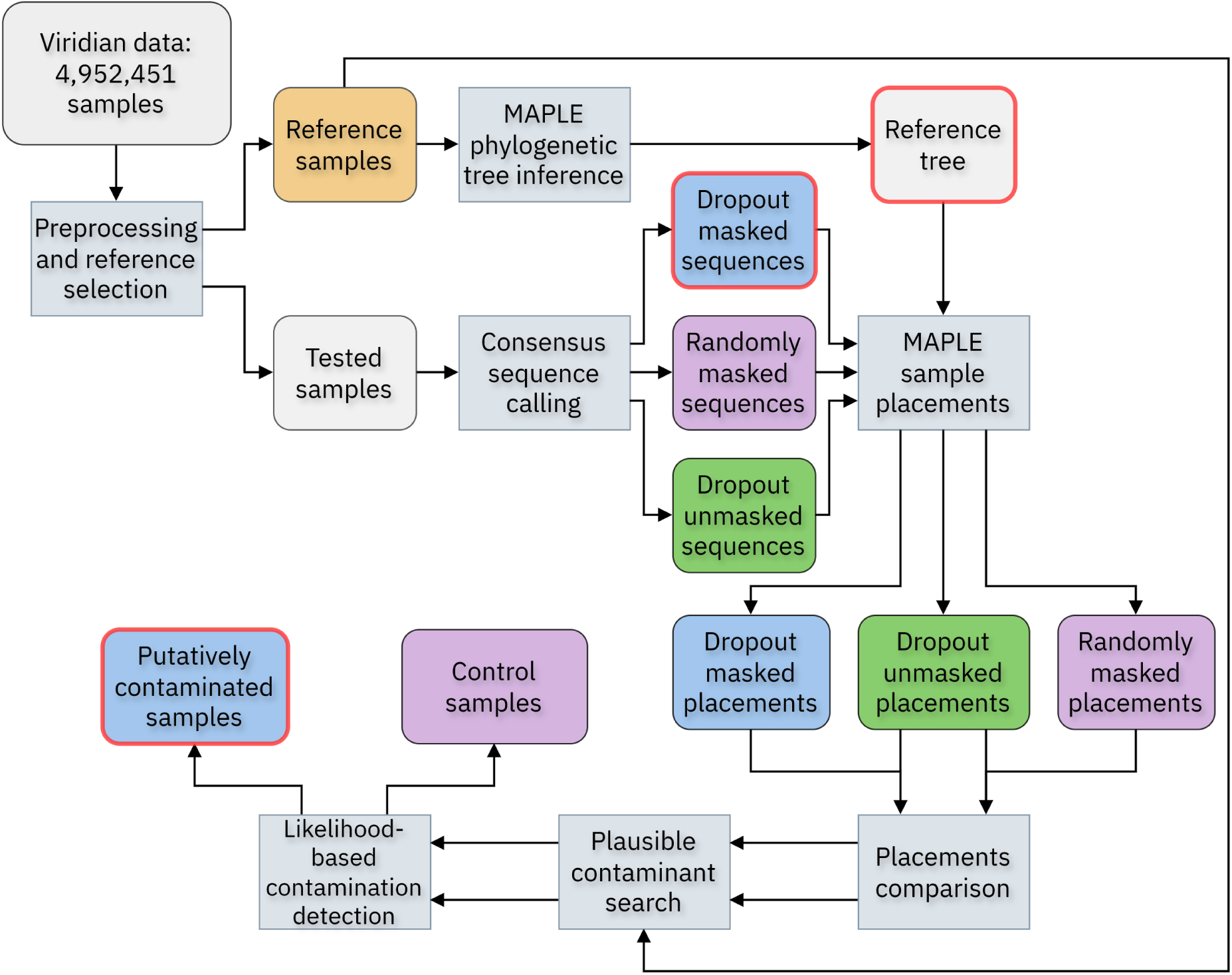
Summary of our PhyCD pipeline. Rounded-edge blocks correspond to data, while sharp-edged blocks describe computational steps. The three blocks outlined in red are the main outputs of the pipeline, accessible on our Zenodo repository (see Data availability). From the top left, we start by selecting samples with few “dropout-masked” positions (see Section 2.2) and heterozygous sites. These 785,011 samples constitute a “reference” set (orange block) used to build a reference phylogenetic tree, and do not undergo the rest of the pipeline (see Section 2.3.1). We then dropout-mask the remaining “tested samples” to address possible combinations of amplicon dropout and low-proportion contamination (see Section 2.3.2 and Figure 1); we call the resulting consensus sequences “dropout masked” (highlighted in blue). We also retain “dropout unmasked” sequences (highlighted in green) without dropout masking (but other masking steps during preprocessing still apply) for later use. We also generate “randomly masked” genomes (highlighted in purple) by replacing dropout masking with a random masking of the same number of nucleotides for masking performance evaluation (see Section 2.3.2). Dropout masked, dropout unmasked, and randomly masked sequences are then phylogenetically placed onto the reference tree to find their closest ancestor. A dropout masked sequence separated from the tree by substantially fewer nucleotide substitutions than the corresponding dropout unmasked sequence is considered a “putatively contaminated sample” (Section 2.3.3) and is further investigated. For each such suspected contamination, we look for reference samples that are “plausible contaminants” (purple genomes in Figure 1). A plausible contaminant should match the dropout unmasked sequence at dropout masked positions, and should match the minor alleles at heterozygous sites elsewhere. This match is assessed statistically to infer “corroborated contaminations” (see Section 2.3.4).

#### 2.3.1 Reference dataset and reference phylogeny

We select a subset of “reference” samples for which contamination is unlikely, or at least unlikely to have impacted the consensus sequence of the sample. We use these samples to infer a reference phylogenetic tree that will serve as a reliable backbone on which all other (“tested”) samples will be placed to highlight potential consensus sequence errors due to contamination and amplicon dropout (described in the following sections).

For the selection of these reference samples, we assume that samples with few heterozygous sites and without large regions of substantial coverage drop are unlikely to have a consensus sequence affected by contamination. Contamination with two similar viral lineages, even at a high pro-portion of the contaminant genome within the sample, is unlikely to lead to consensus sequence calling issues and so from our perspective it can be ignored. The first requirement for samples to have few heterozygous sites is motivated by the assumption that the contamination with a substantially different lineage from the primary one in the sample will typically cause heterozygosity in the read data at genome positions in which the primary and the contaminant genomes differ. It is common however for samples to have a small number of heterozygous sites, for example due to within-host mutations, and so our heterozygosity threshold cannot be too stringent. This requirement might how-ever not be sufficient to detect contamination if the contaminant is present at very low proportion compared to the primary genome. The second requirement of not having large drops in coverage is motivated by the fact that in a sample contaminated with a low-proportion contaminant (which might therefore not trigger the first filter), an amplicon dropout in the major genome, possibly detectable as a large coverage drop, might still cause issues with consensus sequence calling (Figure 1), which we want to avoid in our reference tree.

In summary, we will include a sample *s* in the reference set if:

- it has *κ* or fewer positions under the coverage threshold described in Section 2.2 *ρM_s_*, with *M_s_* the median coverage of *s* and *ρ* our frequency threshold;
- it has *θ* or fewer heterozygous sites, a heterozygous site being a position with minor allele proportion over *η*.

We tried several combinations of parameters (*ρ*, *κ*, *η*, *θ*). Ultimately, the target parameter values should be strict enough (lower values of *κ*, *θ* and *η*; higher values of *ρ*) to remove most samples possibly affected by contamination-related consensus sequence issues, but not too strict as to filter out too many samples and end up with a phylogenetic tree not sufficiently representative of the total true evolutionary history of the virus. The final default parameters selected for the pipeline are (*ρ* = 5%, *κ* = 0, *η* = 10%, *θ* = 3), resulting in 785,011 reference samples (Figure 3A–B). The reference phylogeny was inferred with the pandemic-scale phylogenetic inference tool MAPLE v0.7.5 with options “--model UNREST --rateVariation”. Despite including only a minority (15.9%) of the sample, the reference tree includes the majority of Pango lineages, particularly those represented by many samples (Figure 3C).

**Figure 3:**
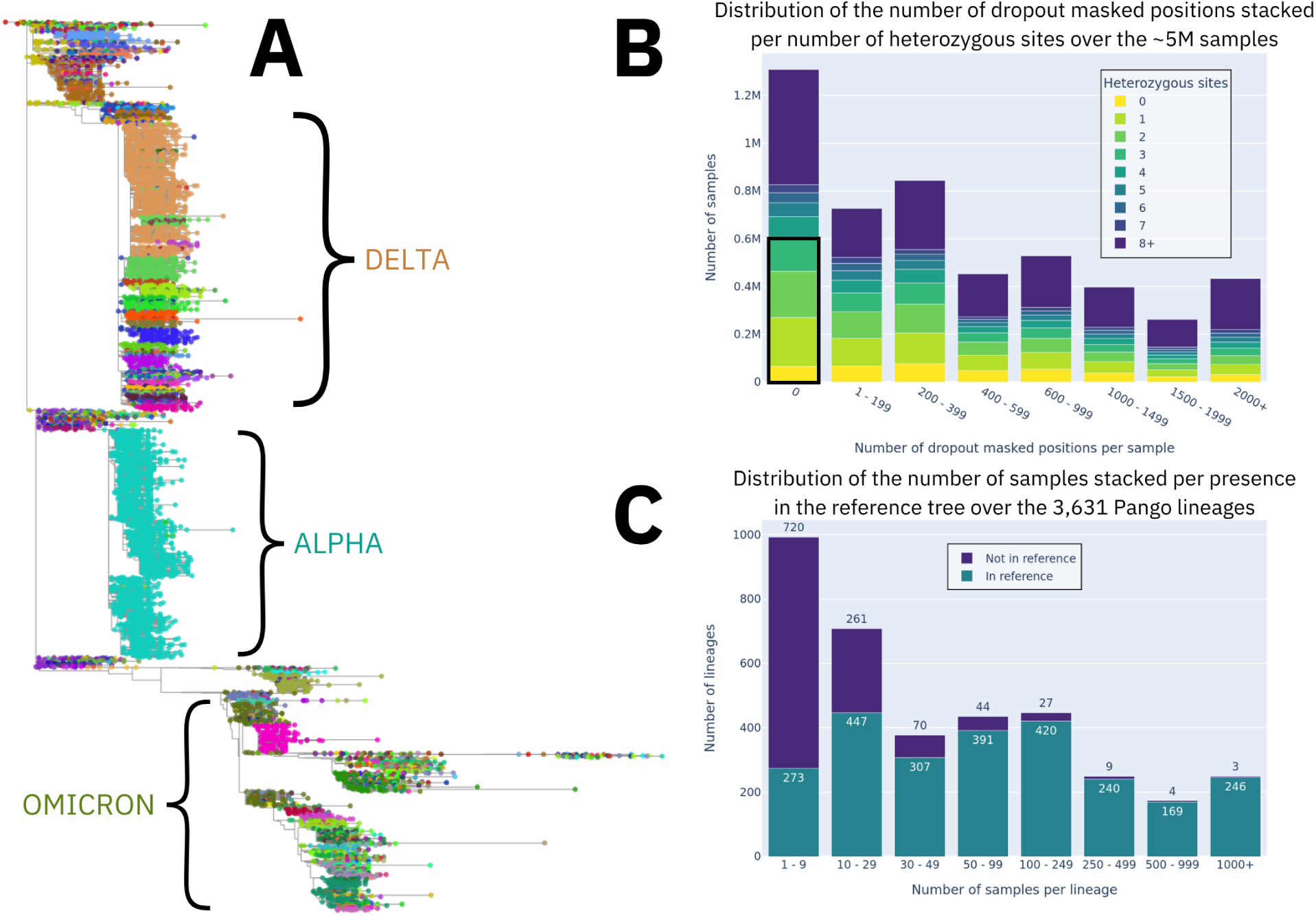
**A,** Reference phylogenetic tree, visualized with Taxonium [23], built with 785,011 reference Viridian samples [11] (Section 2.3.1). Each terminal node in the tree (shown as a coloured circle) corresponds to a sample, with colour indicating its Pango lineage [20] assigned by Pangolin [17]. The names of the main SARS-CoV-2 variants are displayed alongside the corresponding regions of the tree. **B,** Stacked barplot showing the distribution of number of “dropout masked” positions per sample (*x*-axis), over all the 4,952,451 samples. Dropout masked positions are those under the
*pM_s_* coverage threshold (see Section 2.3.1) with *ρ* = 5%. The numbers are stacked per number of heterozygous sites with minor allele proportion over *η* = 10%. A black rectangle shows the samples retained to build the reference tree with default parameter values *K* = 0 and *θ* = 3. **C**, Stacked barplot showing the distribution of the number of samples of Pango lineages. These are stacked according to their presence or absence in the reference tree. Out of 3,631 lineages, 2,661 are present in the reference tree, and of the 970 missing lineages only 25 include more than 100 samples.

#### 2.3.2 Consensus sequence calling

The consensus sequence calling step outputs three sequences per sample: one masked according to the dropout masking principle described in Section 2.2 (“dropout masked”); one with the same number of genome positions masked, but randomly chosen (“randomly masked”), which we use later in the pipeline as a control; and one without any dropout masking applied (“dropout unmasked”) which we use to benchmark the impact of our masking. The latter sequence can be considered as the sequence that would typically be used for downstream analyses when not using our dropout masking approach. Note that the preprocessing steps mentioned in Section 2.1 still apply to this sequence. These three sequences are generated for each tested sample (those not selected as reference in the previous section).

We generate the randomly masked consensus sequences by shifting the dropout masks by a random number of nucleotides between 1,000 and 29,000 (wrapping back to the start of the genome when the shifted masking position exceeds the genome length). In case of overlap between our randomly shifted masking and previous maskings applied during the preprocessing step (Section 2.1), we further mask genome positions until an equal number of positions is masked in our “dropout masked” and “randomly masked” sequences.

#### 2.3.3 Phylogenetic placements

For each tested sample (4,167,440 out of 4,952,451 total, i.e. 84.1%), the three corresponding consensus sequences (dropout masked, dropout unmasked, and randomly masked) are then phylogenetically placed onto our reference phylogeny (Section 2.3.1) using MAPLE v0.7.5 [5–7] with options “--model UNREST --rateVariation” and “--minBranchSupport 0.005”. Phylogenetic placement identifies the evolutionarily closest genome in the reference tree to each placed sequence. The phylogenetic distance (branch length) separating the placed sequence from the reference tree rep- resents the number of substitutions separating the placed sequence from the reference dataset, and, as we assume that the reference sequences are not contaminated, these substitutions might have been caused by private mutations along the lineage leading to the sampled genome, or by consensus sequence errors potentially caused by contamination. A random masking strategy is equally likely to mask private mutations and sequence errors. Our dropout masking is instead aimed at removing potential sequence errors, and it should be agnostic towards private mutations; therefore, assuming it is effective, we expect it on average to reduce the placement branch length more than the random masking strategy, and roughly by the additional number of consensus sequence errors prevented compared to a random masking strategy. Unmasked sequences should instead always have the longest placement branch length.

The pandemic-scale phylogenetic approach of MAPLE allows us to perform millions of phylogenetic placements on a large reference tree. Placements are done using the MAPLE option “--findSa-mplePlacement”, which we specifically implemented to search for the optimal placement of the considered sequence onto the reference phylogenetic tree without actually adding the sequence to the tree.

For each placed sequence, MAPLE summarizes:

- the most probable placement branch in the tree;
- alternative plausible placements and their support [6] (see e.g. Figure 4);
- the evolutionary distance (“branch length”) between the considered sequence and the tree, that is, the number of substitutions separating the sequence from the most likely ancestor genome in the tree;
- the list of substitutions separating the considered sequence from the most likely placement in the tree.

**Figure 4:**
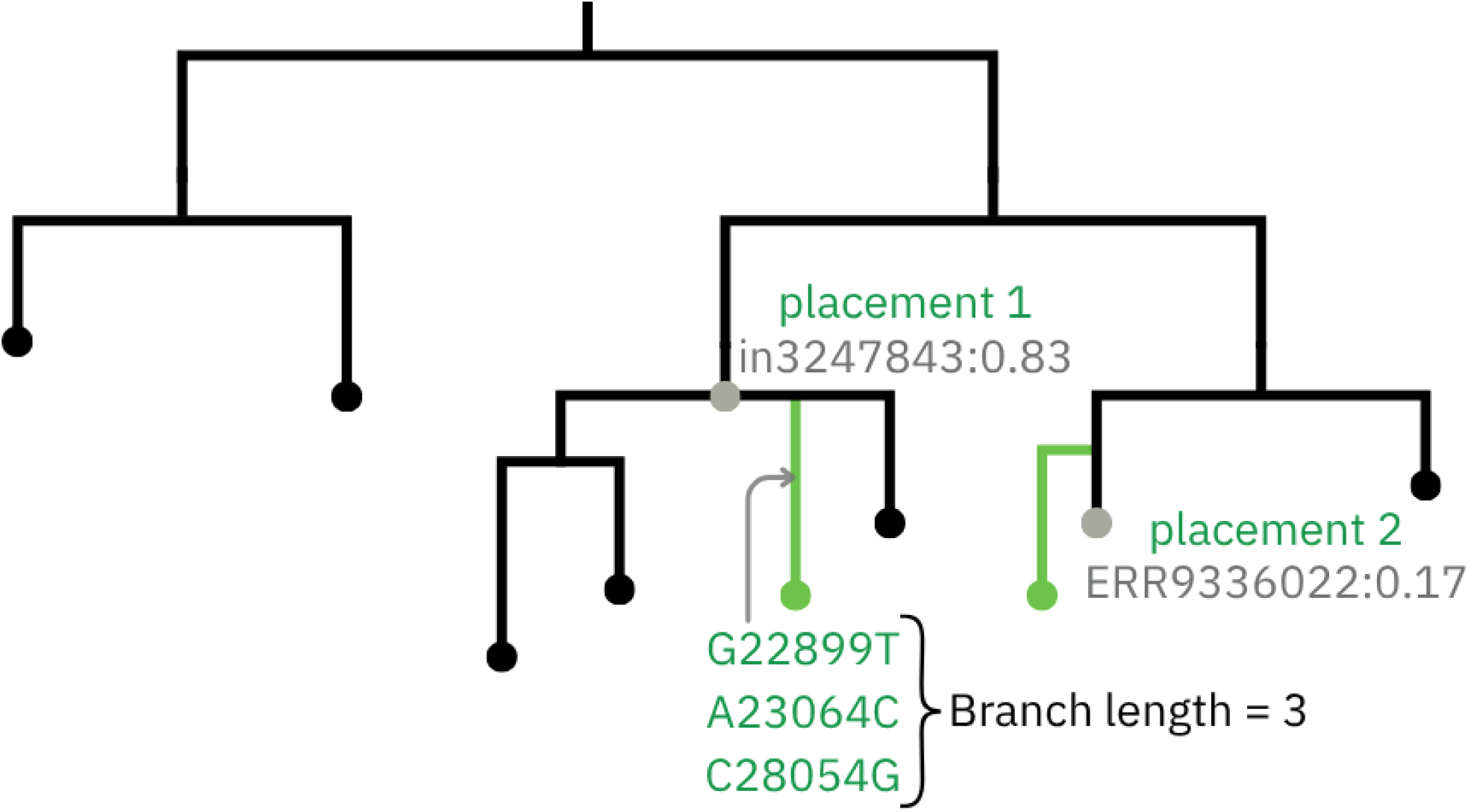
Example phylogenetic sequence placement outcome. We show possible placements (green) of the considered sequence onto the reference phylogeny (black). These two mutually exclusive placements have support probabilities respectively of 0.83 and 0.17. For the most likely placement we also show an example list of substitutions separating the sequence from the tree. “G22899T” for example means a substitution from nucleotide G to nucleotide T at genome position 22899. The placement branch length, equal to 3 in this case, is the number of substitutions separating the sequence from its placement node.

We expect that a combination of contamination and amplicon dropout might introduce errors in a sample consensus sequence, and that these errors would usually not be matched by actual mutations represented in our reference phylogeny within the same genetic background of the genome being sequenced. Therefore, we expect that a consensus sequence containing such errors would often have higher distance from the reference tree than the same sequence in which these errors had been masked (Figure 5). Following this principle, we calculate the difference in placement branch length between the dropout masked and the dropout unmasked sequence of the same sample as a measure of the effectiveness of our dropout masking. To assess the significance of these difference (we expect that some private mutations or consensus sequence errors can be masked in the dropout masking sequence just by chance), we also calculate the differences in placement branch length between the randomly masked sequences and the dropout unmasked sequence.

**Figure 5:**
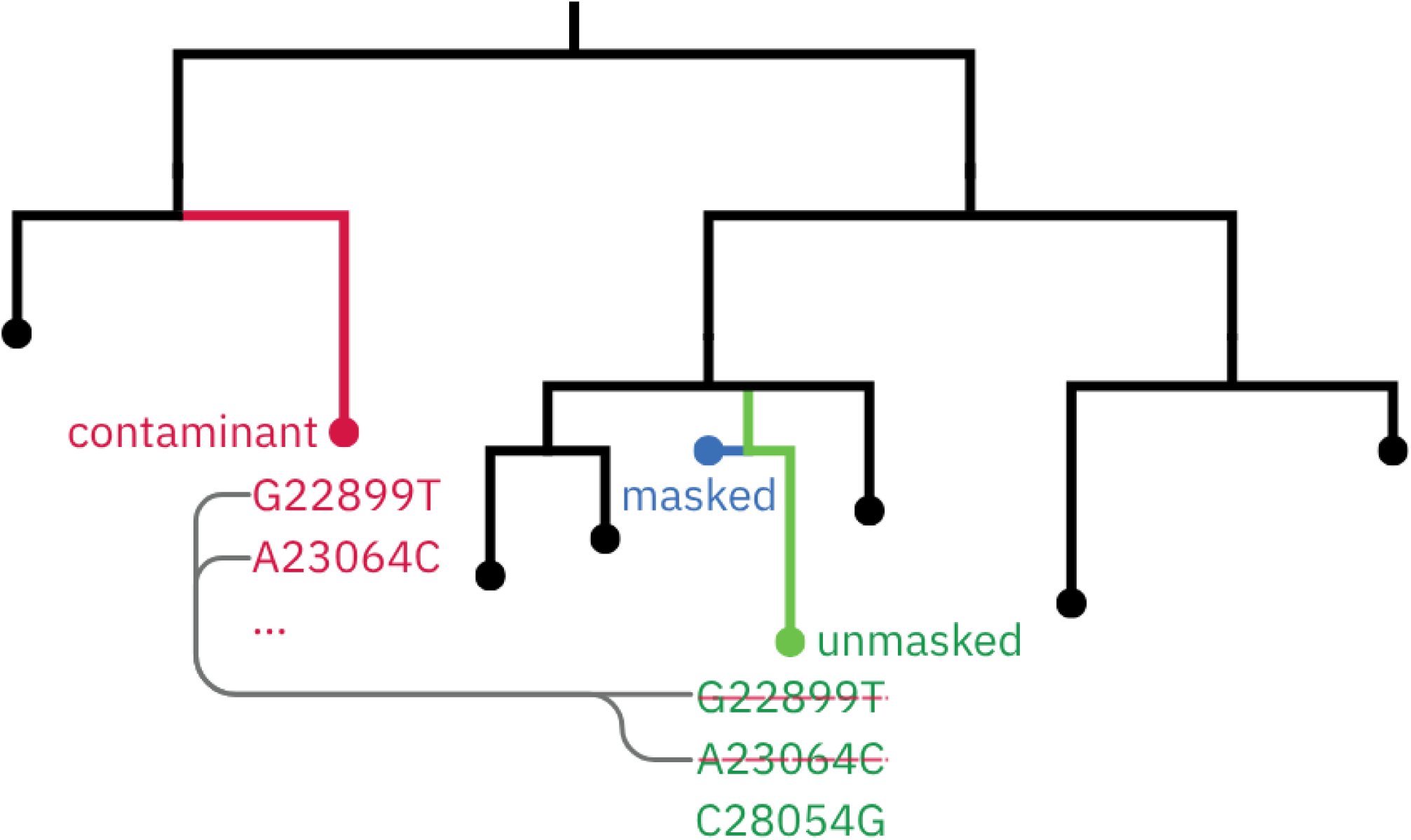
Example ideal scenario of the dropout masking successfully removing substitutions caused by contamination. For a single considered sample, we show in green the placement of the “dropout unmasked” consensus sequence, in blue the placement of the “dropout masked” one, and in red a candidate contaminant for this sample. Here the dropout masked sequence has a shorter placement branch length than the unmasked one, suggesting that the masking might have prevented two consensus sequence errors (those struck with a red line, while C28054G is present in the dropout masked sequence as well). These two prevented substitutions are found also in another genome within the reference phylogeny, so this genome (red) is considered a plausible contaminant for the considered sample.

Next, we want to select a set of samples that are likely to have been affected by contamination, and for which PhyCD is likely to have prevented errors in the consensus sequence. We select as putative contaminated samples those where the dropout masked sequence is 2 or more substitutions shorter than the corresponding dropout unmasked sequence placement branch length (meaning that the dropout masking putatively prevented at least 2 consensus sequence errors). From these samples we filter out those where the dropout masked sequence has a phylogenetic placement branch length over 5: such a high distance is rare and might be caused by long-term within-host evolution. In total, with these thresholds we identify 10,942 putative contaminated samples. Applying the same procedure to the randomly masked sequences we find just 5,457 samples. This suggests that our dropout masking prevents errors in more than 5,000 consensus sequences. However, it also indicates that for thousands of samples our dropout masking might remove genuine genetic information. We try to reduce this number of false positives by further refining our selection of putative contaminated samples by assessing candidate contaminants, as described below.

#### 2.3.4 Contaminant search and likelihood-based assessment

We explored whether putative contamination events could be corroborated on a per-sample basis by identifying plausible contaminant genomes among the reference samples. For each of the 10,942 candidate contaminated samples, we searched for candidate contaminants among the 785,011 reference genomes using two complementary criteria: first, similarity to the dropout unmasked consensus sequence within the dropout masked regions (the regions where a contaminant’s reads would be expected to dominate); second, similarity to the minor alleles observed at heterozygous sites across the rest of the genome (where the contaminant would be present only at low proportion). These two criteria were combined into a single closeness score to rank candidate contaminants for each sample (see Supplementary Section S1.1). Considering that the search is only done among the reference genomes, not finding a plausible contaminant for a flagged sample does not necessarily rule out the contamination hypothesis, but it does decrease its support.

Candidate contaminants were then also probabilistically assessed using a likelihood-based model adapted from Eyre et al. [8]. The model computes the likelihood of the observed read-level allele counts assuming that the reads derive from a mixture of the primary genome and the candidate contaminant (see Supplementary Section S1.2).

Unfortunately, neither approach achieved useful discriminatory power (see Supplementary Section S1.3), so while these two steps are included as an optional part of the PhyCD pipeline, they were not used in the further filtering the 10,942 candidate contaminated samples investigated in the Results section.

## 3 Results

### 3.1 Masking and contamination identification performance

As described above, our pipeline identifies and masks 10,942 putative contaminated samples out of a total of 4,952,451 SARS-CoV-2 sequencing samples. However, not all of these are expected to be actual contaminations. Our pipeline in fact highlights 5,457 samples that had previously been randomly masked (Figure 6A,B). These 5,457 samples give us a rough idea of how many contamination false positives we expect from our pipeline, since by masking regions of the consensus sequences we are bound to also mask genuine private mutations in our samples. While only around half of the identified 10,942 putative contaminated samples are therefore expected to be actual contaminated samples with errors in their consensus sequences, it can be argued that masking all these 10,942 samples, and therefore losing some genuine mutational information, is a good trade-off if it prevents errors and biases from downstream analyses. As further evidence of the authenticity of some of the identified putative contaminated samples, we also find that dropout masking removes more substitutions than random masking (Figure 6C,D), while requiring fewer masked positions per prevented substitution (Figure 6E,F).

**Figure 6:**
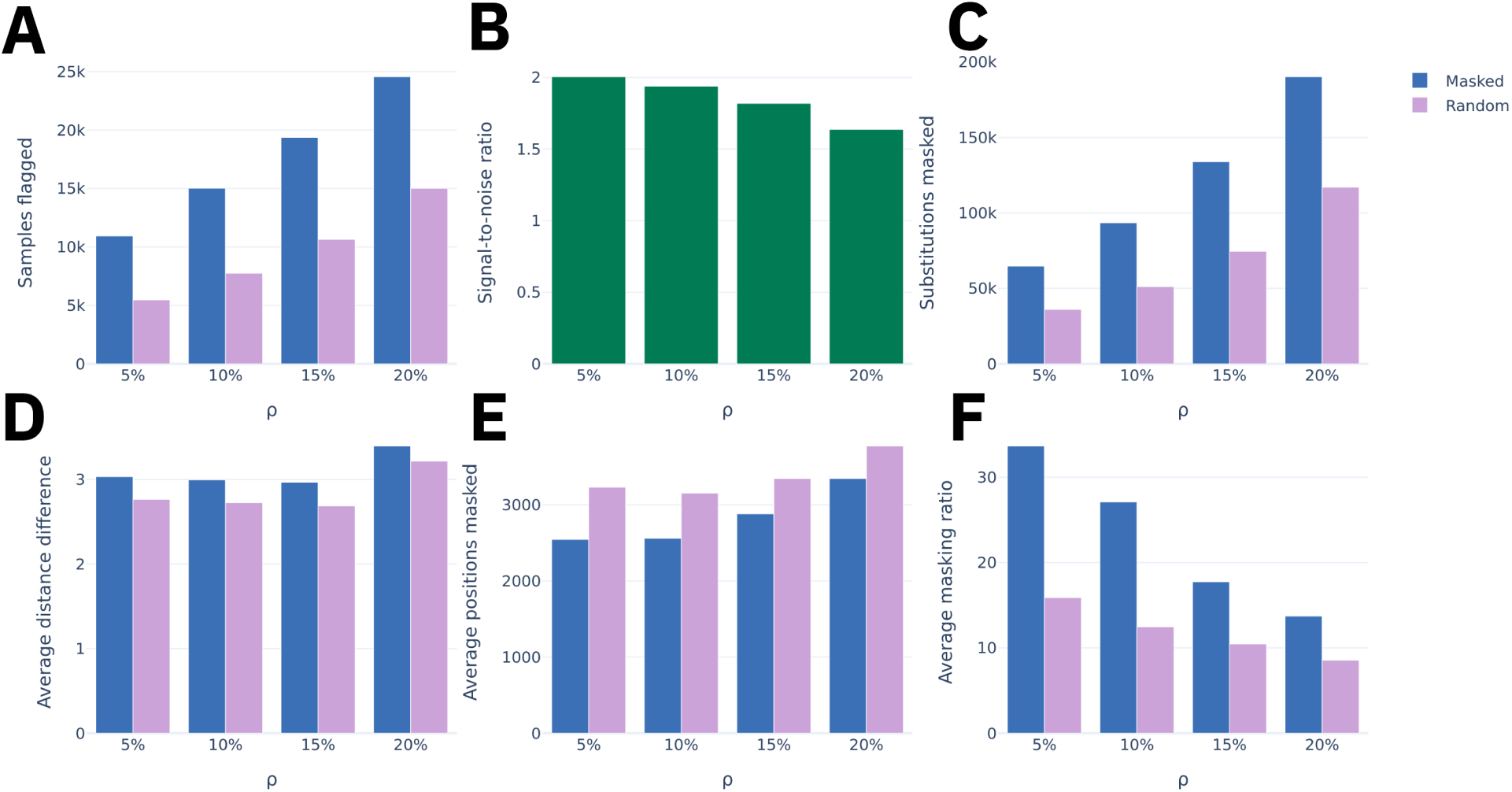
Performance evaluation of our pipeline PhyCD. On the *x*-axis of all panels we have the four tested values of *ρ* (5%, 10%, 15%, and 20%). Blue bars refer to the dropout masked sequences and purple bars to the randomly masked control baseline. **A,** Total number of flagged putative contamnated samples. **B,** Signal-to-noise ratio, defined as the number of flagged samples in the dropout masked dataset divided by the number in the randomly masked dataset. **C,** Total number of mutations removed by the masking from phylogenetic placement branch lengths. **D,** Average distance difference, showing the mean reduction in placement branch length between the unmasked and masked consensus sequences among flagged samples. **E,** Average number of genome positions masked per flagged sample. **F,** Average masking ratio, defined as the proportion of distance reduced over the proportion of the genome masked, indicating the efficiency of distance reduction per masked nucleotide.

These results were obtained with *ρ* = 5%, where *ρ* is to the proportion of the median coverage under which positions are masked. Increasing *ρ* leads to an increase in the number of masked positions. To evaluate the impact of this threshold, we compared pipeline outcomes across values of *ρ* at 5%, 10%, 15%, and 20% (Figure 6). Values of *ρ* substantially under 5% do not lead to different outcomes as the strict 50X threshold then always overrides the adaptive threshold.

We find that increasing *ρ* results in more putatively contaminated samples being flagged by the pipeline but also in more samples flagged in the randomly masked control dataset (Figure 6A). The signal-to-noise ratio decreases with higher *ρ* (Figure 6B), from 2.0 at *ρ* = 5% to approximately 1.6 at *ρ* = 20%. This suggests that while a higher *ρ* can address more potential issues, it also introduces proportionally more false positives and therefore unnecessary losses of genuine mutational information. It is worth noting that, since the number of flagged samples in the randomly masked dataset is considered approximately equal to the number of false positives in the dropout masked dataset, then the signal-to-noise ratio is the inverse of the false positive rate.

The total number of masked substitutions also increases with *ρ*, but so does the number masked substitutions in the random control (Figure 6C,D).

As *ρ* increases, the average number of genome positions masked per flagged sample rises substantially (Figure 6E), from approximately 2,500 at *ρ* = 5% to over 3,500 at *ρ* = 20%, leading to a steep decline in the number of substitutions prevented per base masked (Figure 6F).

Taken together, these results highlight the success in masking substitutions attributable to contamination, but also the difficulty in pinpointing contaminated samples with high confidence. While a more aggressive masking strategy (higher *ρ*) removes more errors from putative contaminations, it however degrades both the signal-to-noise ratio and the overall efficiency of the masking. The optimal choice of *ρ* probably depends on the context, that is if one wants to prioritise sequence data error removal or the mutational informativeness of the data. In practice, we recommend a conservative setting (*ρ* = 5%) to maximize the signal-to-noise ratio.

In these analyses we used fixed values *κ* = 0, *η* = 10%, and *θ* = 3. These parameters only impact the selection of reference samples, not their masking, and were chosen to obtain a clean but representative reference tree (Figure 3).

We also investigated if other summary statistics could help distinguish genuinely contaminated samples from false positives, so to further refine our set of putative contaminated samples. We designed and evaluated a series of such contamination metrics, described in Supplementary Section S1.3. However, we find that these summary statistics lack discriminatory power to improve the identification of true contamination events (Figure S1).

### 3.2 Analysis of an example putative contaminated sample

To showcase how we can investigate putatively contaminated samples in further detail, we present an in-depth analysis of an example sample, SRR24221569, taken among the 10,942 putative contaminated samples selected in Section 2.3.3.

Sample SRR24221569 has two unique contaminant candidates with the same score. For definitions of these contaminant scores, see Supplementary Section S1.1 and S1.2; in short, this means that two separate reference genomes could explain a substantial portion of dropout masked substitutions and within-sample heterozygosity, and therefore both genomes are good candidates for being the minority contaminating variant within sample SRR24221569.

According to the phylogenetic placements of the sample, the dropout masking reduced the placement branch length from 2.1 to 0, and masked 566 positions (approximately 2% of the genome). Among those positions we masked 7 mutations (with respect to the reference): C4423T, G22992A - A23013C - T23018G - A23055G - T23075C, G29734C (mutations are grouped by proximity). Masking increased the placement support from 0.32 for the unmasked sequence to 0.94 for the dropout masked sequence; contamination-related sequence errors can result in apparent recombinant sequences, and these recombinants can have uncertain phylogenetic placement, so the increase in placement confidence observed could be a result of the removal of contamination-related sequence errors.

Figure 7 displays the read data at the masked block between positions 22850 and 23150 of the reference sequence. As expected, the coverage at the masked positions is over 20X (the Viridian threshold), but is under our threshold, at around 2.6% of the sample median coverage. This suggests a possible combination of amplicon dropouts and contamination at these positions, successfully detected by our approach. Furthermore, heterozygosity at position 22882 (just outside the masked block) shows that reads covering the masked region share the allele 22882G while most other reads have allele 22882T. This strongly supports the claim that those two sets of reads come from different genomes and the hypothesis of the sample being contaminated (or representing a mixed infection).

**Figure 7:**
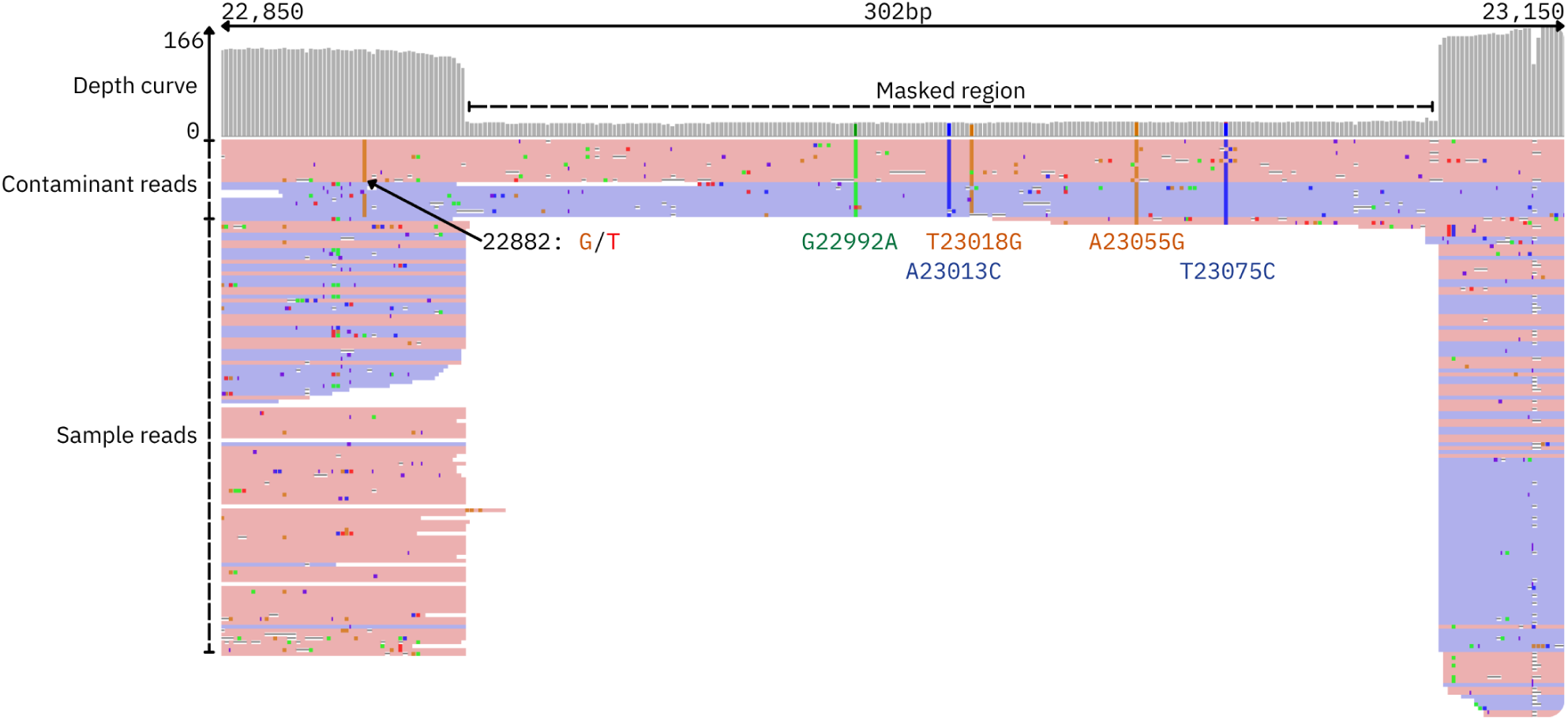
Read data near a masked genomic region of sample SRR24221569 between positions 22850 and 23150 of the reference sequence, visualized using IGV [21]. This sample has a median coverage of 951X. Coverage is around 25X in the masked region and around 160X in the surrounding regions. Masked mutations G22992A, A23013C, T23018G, A23055G and T23075C are shown. Heterozygosity at position 22882 between G and T shows a discrepancy between the reads covering the masked region (putatively sequenced from the contaminant) and the reads nearby, which are putatively mostly from the main genome in the sample.

Figure 8 shows the phylogenetic placements of the sample before and after masking, and the phylogenetic location of the two candidate contaminants. The sample is assigned to lineage BA.5.2.1 both before and after masking. Before masking, its placement branch is 2.1 substitutions long, indicating that no sample in the reference phylogeny is identical to it. After masking, it is placed directly onto an existing sample (SRR21269384) with a placement branch of length zero. Both putative contaminant genomes are identical to the considered sample SRR24221569 within the putative dropout region 22992–23075, where they have high depth (>800X), and they also have mutation T22882G which appears as a heterozygosity within SRR24221569. However, among the two other masked mutations on the placement branch of SRR24221569, C4423T is only present in candidate contaminant sample SRR21561497, while G29734C is only present in the candidate contaminant sample SRR21761600. This means that neither candidate alone accounts for all mutations within the masked regions, and therefore neither is a perfect explanation for the observed data.

**Figure 8:**
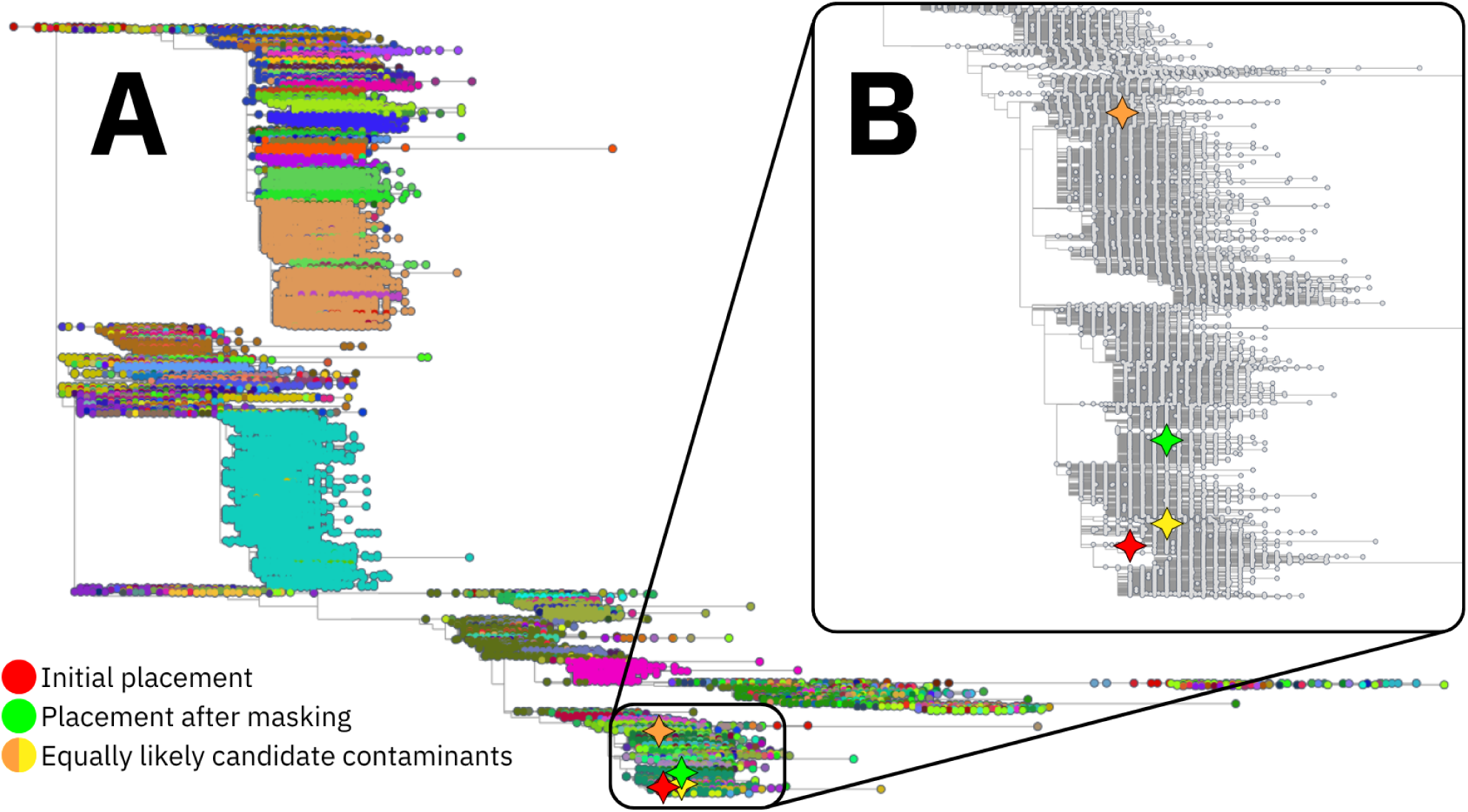
**A,** Reference tree as shown in Figure 3, coloured by Pango lineages [17], visualized with Taxonium [23]. Four positions are highlighted by large stars: the initial placement of the putative contaminated sample SRR24221569 on the reference tree (red star), its placement after masking (green star), and the two best candidate contaminants (SRR21561497 in orange, found in lineage BA.5.5, and SRR21761600 in yellow, found in lineage BF.21). **B,** Zoom-in on the BA.4 lineage (Omicron variant).

Sample SRR24221569 is not exceptional among the flagged samples: its distance reduction of 2.1 is close to the dataset average of 3.0 among the “dropout masked” flagged samples. While this in-depth level of manual investigation of the read data and phylogenetic patterns could be more generally useful, it cannot currently be performed on all 10,942 flagged samples.

## 4 Discussion

In this study we presented PhyCD, a phylogeny-aided computational approach for investigating consensus sequence errors caused by the combination of sample contamination and amplicon dropout. By masking genome regions exhibiting substantial drops in sequencing coverage and evaluating the resulting sequences via phylogenetic placement on a pandemic-scale reference tree, we aimed to separate sequencing artifacts from genuine viral evolution. To rigorously assess the performance of our consensus sequence masking approach, we benchmarked it against a randomly masked control dataset.

Applying PhyCD to nearly 5 million SARS-CoV-2 genomes, we identified 10,942 putatively contaminated samples, against 5,457 found from random masking, suggesting that around 50% of these putatively contaminated samples might be false positives. We unsuccessfully attempted to reduce the number of expected false positives by using various summary statistics to identify true contaminated samples.

While a 50% rate of false positives might be considered high in some settings, in this scenario losing (or partially masking) around 5,000 non-contaminated samples to be able to account for around 5,000 contaminated samples should be considered very advantageous. In fact, these samples were selected for the potential impact of contamination on their consensus sequence, and therefore on downstream analyses. Analyses likely to be adversely affected by contaminated sequences include the inference of recombination, transmission, phylogenetic tree inference, lineage assignments, and many others. Also, removing or partially masking around 5,000 genomes from a total of 4,952,451 (0.1%), is likely to lead to only very limited loss of true evolutionary signal.

Our investigation highlighted the fundamental methodological trade-off between sensitivity and precision. While more aggressive masking strategies (e.g., *ρ* = 20%) flag a higher absolute number of contaminations, they severely degrade the signal-to-noise ratio and masking efficiency, introducing unnecessary maskings to the consensus genomes. This compromise is not easily resolved in the contamination detection task due to the lack of data needed to get higher precision. By characterizing this trade-off, we show that the balance between sensitivity and specificity in coverage-based contamination detection is context-dependent. One could imagine that in global-scale surveillance databases where maintaining a strictly contamination-free cohort is paramount, researchers might opt for a high *ρ* threshold followed by the strict exclusion of all flagged samples. Conversely, for smaller, localized datasets where maximizing data retention could be critical, users could apply a conservative low *ρ* threshold and utilize dropout masked sequences in downstream evolutionary analyses instead of completely removing putative contaminated genomes.

While PhyCD was developed for and applied to SARS-CoV-2, it could in principle be extended to other amplicon-sequenced pathogens and could be useful in potential future pandemics. However, two key prerequisites determine its applicability. First, PhyCD requires a large and sufficiently dense set of reliable genomes to inform an encompassing backbone reference phylogeny. Second, for a pathogen with much higher mutation rate and sparser sampling than SARS-CoV-2, each genome would naturally accumulate multiple mutations relative to the closest relative in the reference tree, and the additional substitutions introduced by contamination would become harder to distinguish from genuine divergence.

Among the limitations of PhyCD, we acknowledge the lack of a comprehensive “gold-standard” dataset of experimentally verified contaminated samples that could help validate true-positive and false-positive rates. A second limitation is that our approach is based on several individual filtering steps, rather than a single analysis encompassing all data considered (phylogenetic placement, read coverage variation, heterozygosity, possible contaminant genomes) within a coherent probabilistic model of contamination. Such a framework might in theory achieve higher power to identify and address contamination, in particular if additional data like the time and location of collection of samples, the location of primers along the genome (as used by Viridian [11]), and the linkage of nucleotide observations within the read data (as for example in Figure 7 where the allele 22882G mostly appears on putatively contaminating reads), would also be integrated. An important challenge of such a framework would however be to achieve high computational performance, that is, to be scalable to millions of sequencing datasets.

While detectable contamination affecting consensus sequences seems to be relatively rare (affecting roughly 0.1% of our dataset based on our inference), the massive scale of SARS-CoV-2 sequencing means that thousands of genomes are likely affected. Every one of these genomes, if left uncorrected, might then cause an artifact, for example in the inference of recombination and new lineages. Approaches like PhyCD, leveraging massive-scale genome data and phylogenies to corroborate read-level anomalies, represent a promising direction for scalably and informatively automating the curation of genomic epidemiological data.

## Data availability

A Zenodo repository is publicly available at zenodo.org/records/19815950. It contains the reference tree presented in Figure 3, a MAPLE format alignment of the 4,952,451 samples with dropout masked positions and the list of the 10,942 putatively contaminated samples. These files were generated using PhyCD with default parameters.

## Code availability

PhyCD code is accessible on GitHub at github.com/oanoufa/PhyCD under the GNU General Public License v3.0. The pipeline was built using Snakemake [16].

## Supplementary sections

### S1.1 Contaminant search

We explored whether individual putative contamination events could be corroborated by identifying plausible contaminant genomes. If for a given putatively contaminated sample no plausible contaminant genome is found in our dataset, then it can be argued that the sample is less likely to be actually contaminated, and the substitutions observed in the “dropout masked” regions of the genome might be real mutations. To identify plausible contaminants, for each of the 10,942 candidate contaminated samples we search among the reference genomes (Section 2.3.1) for those that best match the dropout unmasked consensus sequence within the dropout masked regions (score *s*_1_), and those that best match the minor alleles at heterozygous sites across the whole genome (score *s*_2_).

We define score *s*_1_ as the number of substitutions (with respect to the SARS-CoV-2 reference genome) within the dropout masked regions found both in the “dropout unmasked” consensus sequence of the putative contaminated sample and in the candidate contaminant genome (*n_match_*, see Figure 5), divided by the total number of substitutions within the dropout masked regions in the dropout unmasked consensus sequence of the putative contaminated sample (*n_sample_*) plus the total number of substitutions within the dropout masked regions within the candidate contaminant genome but not in the putative contaminated sample dropout unmasked consensus sequence (*n_extra_*):

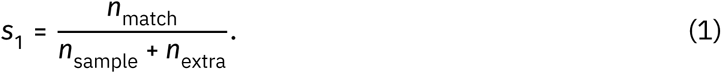

The rationale for *s*_1_ is that candidate contaminants should match the dropout unmasked putative contaminated sample consensus genome at regions of dropout masking (see Figure 1), and this is why *s*_1_ is designed to increase with the number *n*_match_ of substitutions shared between the two sequences in these regions, and decrease with the number of substitutions not shared. As the maximum value of *n*_match_ is bounded by *n_sample_*, *s*_1_ has a maximum value of 1 in case of a perfect match, and a minimum value of 0 in case *n*_match_= 0.

Score *s*_2_ is instead defined as the number of heterozygous sites of the considered putative contaminated sample at which the minor allele (which is therefore not the nucleotide of any consensus sequence for the sample) matches the contaminant candidate sequence (*n_matchHet_*), divided by the total number of heterozygous sites in the considered putative contaminated sample (*n_Het_*):

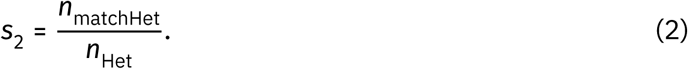

Heterozygous sites are defined as in Section 2.3.1. The rationale of *s*_2_ is that outside of the dropout masked regions the contaminant genotype should be present at a minor frequency (assuming its abundance is high enough to appear in sufficiently many reads outside of dropout regions), appearing as a minor allele in the read data of the putative contaminated sample.

We consider both types of evidence similarly important, so we define a final score *s* (which we call “closeness score”) as the mean of the two component scores:

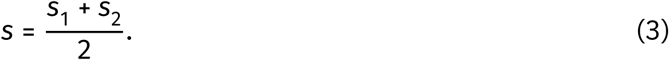

For any considered putative contaminated sample, we track the contaminant candidates with the highest closeness score, which are then probabilistically assessed as described below.

### S1.2 Likelihood-based contamination detection

As a second approach to assess putative contaminated samples and their corresponding candidate contaminants, we use a probabilistic model of contamination. Specifically, we adapt the mixed infection model of Eyre et al. [8]. The Eyre et al. model calculates the likelihood of observed allele counts from pathogen genome sequencing samples assuming that the host is co-infected by two different pre-specified strains of the same pathogen. From the perspective of sequencing data, a mixed infection presents the same patterns as contamination, which justifies the use of a similar model. While Eyre et al. considered a bacterial pathogen, applying the same approach to a virus, as in our case, does not break any of the assumptions of the model.

To reduce computational demand, we only consider sites where the sample or at least one of the candidate contaminants differs from the reference genome. We compare the allele counts from the candidate contaminated sample to each of its candidate contaminants in turn, and calculate a likelihood score for that candidate contaminant and that sample. This likelihood is used to assess how realistic it is that the considered sample has been contaminated by the considered candidate contaminant.

The total likelihood of a pair of a sample and a candidate contaminant is defined as the product of the likelihoods of each individual site. In turn, the likelihood of a site is defined as the product of the likelihoods of all nucleotide observations mapping at the site from all individual reads. More specifically, assuming we have *H* variable sites, that at site 1 ≤ ℎ ≤ *H* we have coverage *X*_ℎ_, that the sample consensus sequence is *S*_ℎ_ at this position, and that the candidate contaminant sequence is *C*_ℎ_, we define the likelihood of the *x*-th read observation *a*_ℎ,*x*_ at site ℎ (with 1 ≤ *x* ≤ *X*_ℎ_) as:

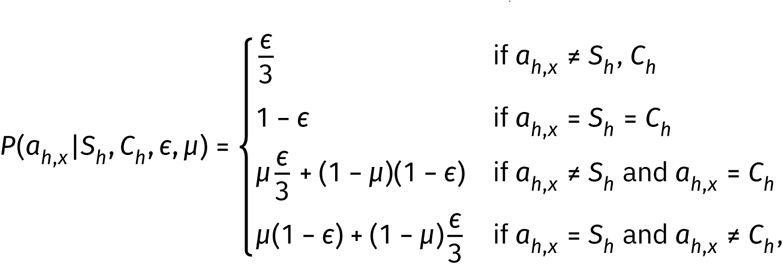

where *ε* is the assumed sequencing error rate, and *μ* is the assumed proportion of the majority genome in the considered sample (and hence 1 − *μ* is the proportion of the contaminant genome in the considered sample). The values of *ε* and *μ* are optimized by maximum likelihood specifically for each genome pair considered within the bounds *ε* ∈ [10^−4^, 0.1] and *μ* ∈ (0.5, 1.0); we record the maximum likelihood value for each sample-contaminant pair (*s*, *c*) as *LK_s_*_,*c*_ = ∏_ℎ_ ∏*_x_ P*(*a*_ℎ,*x*_|*S*_ℎ_, *C*_ℎ_, *ε*, *μ*). Using the same model, we also calculate the maximum likelihood *LK_s_* of sample *s* assuming no contamination; this is done by setting *μ* = 1 (or equivalently assuming that the contaminant is the consensus sequence of *s*, that is *S*_ℎ_ = *C*_ℎ_ ∀ℎ ∈ {1, …, *H*}). We then assume a prior probability of *π*_con_ = 0.001 of a sample being contaminated (loosely informed by the empirical finding of >5000 putatively contaminated samples in our dataset in excess to those expected by chance: Section 2.3.3). We also assume that each contaminant *c* from our candidate selection is equally likely to be a contaminant of any given sample: *π_c_* = 1/*n*_candidates_ with *n*_candidates_ the number of considered candidates for the considered sample. Then, ignoring uncertainty in *μ* and *ε* and possible contaminants outside our set of candidate contaminants, and using Bayes’ theorem, we approximate the posterior probability *P*(*s*_con_) of sample *s* being contaminated as:

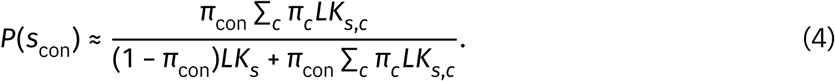

### S1.3 Benchmark of contaminant assessments and summary statistics

Using *ρ* = 5%, we compared the 10,942 “dropout masked” and 5,457 “randomly masked” flagged samples across multiple metrics: number of heterozygous sites (Figure S1A), masking ratio (Section 3.1 and Figure S1B), phylogenetic placement support (Section 2.3.3 and Figure S1C), number of candidate contaminants (Figure S1D), closeness score (Supplementary Section S1.1 and Figure S1E), and contamination posterior probability (Supplementary Section S1.2 and Figure S1F). Our aim is to find ways to distinguish these two sets of samples and therefore improve our ability to identify contaminated samples. All metrics except the masking ratio however show near-total overlap between the two groups, indicating that they lack discriminatory power.

The masking ratio (Figure S1B) shows a partial difference between the two groups: the mean is notably higher for the dropout masked samples than for the randomly masked controls. However, the medians of the two distributions are very close, and the overall distributional overlap is substantial, meaning that the masking ratio alone cannot reliably classify genuinely contaminated samples. Taken together, these results confirm that while our dropout masking performs better than random masking at the aggregate level (see Section 3.1), this aggregate signal does not translate into sufficient per-sample discriminatory power to further refine our set of putative contaminated samples.

**Figure S1:**
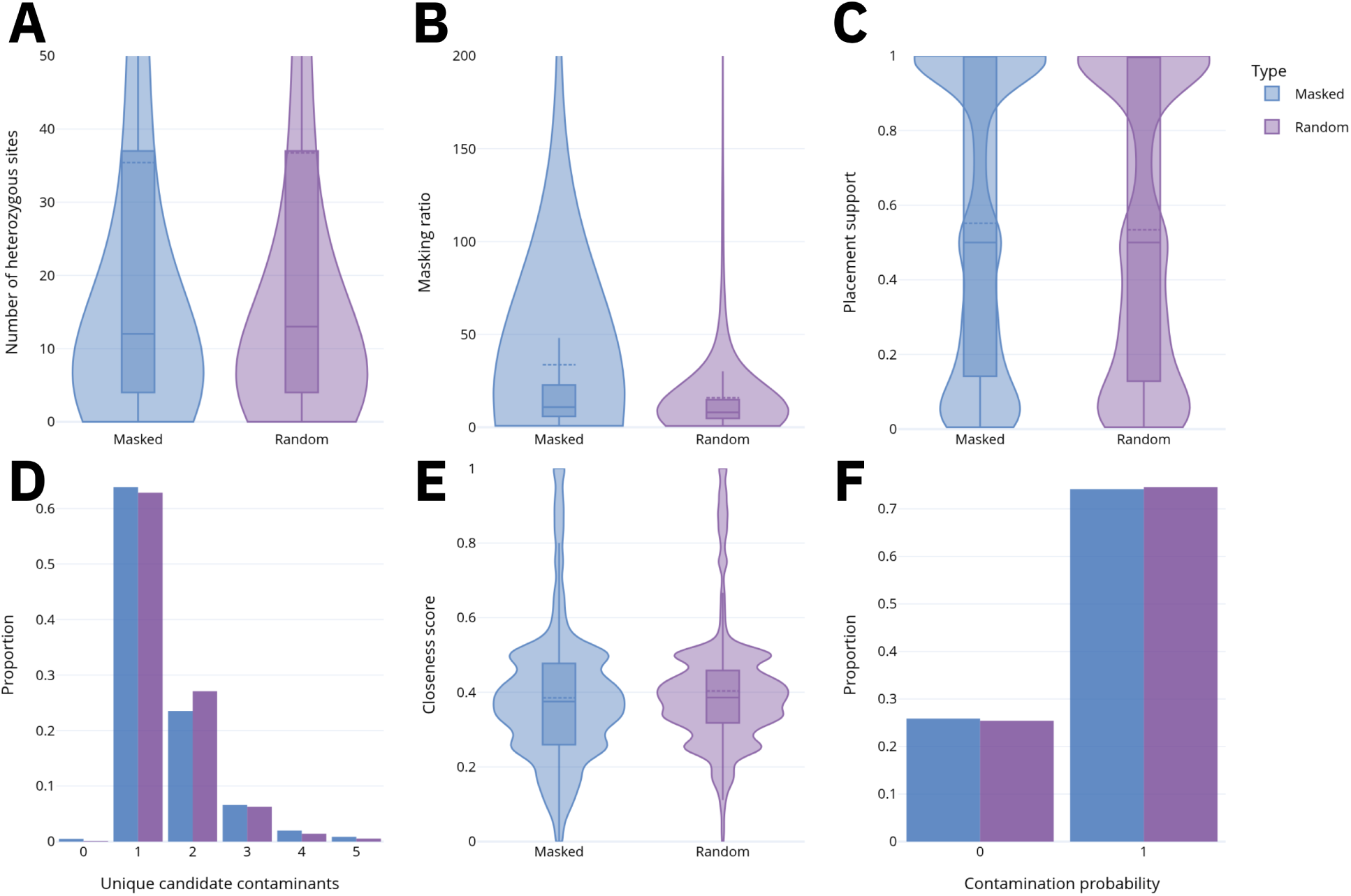
Comparison of evaluation metrics between putatively contaminated samples (blue) and control samples (purple). For all plots we used *ρ* = 5%. **A,** Number of heterozygous sites. **B,** Masking ratio, indicating phylogenetic placement distance reduction per masked nucleotide (Section 3.1) **C,** Phylogenetic placement support (Section 2.3.3). **D,** Number of unique candidate contaminants identified in the reference tree. **E,** Closeness score (s) of the sample’s best candidate contaminant (Supplementary Section S1.1). **F,** Result of the probabilistic test of contamination of Supplementary Section S1.2 (0 = uncontaminated, 1 = contaminated; only posterior probabilities very close to 0 or 1 were encountered).

